# Pooled optical screening in bacteria using chromosomally expressed barcodes

**DOI:** 10.1101/2023.11.17.567382

**Authors:** Ruben R. G. Soares, Daniela A. García-Soriano, Jimmy Larsson, David Fange, Dvir Schirman, Marco Grillo, Anna Knöppel, Beer Chakra Sen, Fabian Svahn, Spartak Zikrin, Mats Nilsson, Johan Elf

## Abstract

Optical pooled screening is an important tool to study dynamic phenotypes for libraries of genetically engineered cells. However, the desired engineering often requires that the barcodes used for *in situ* genotyping are expressed from the chromosome. This has not been possible in bacteria. Here we describe a method for *in situ* genotyping of libraries with genomic barcodes in *Escherichia. coli*. The method is applied to measure the intracellular maturation time of 81 red fluorescent proteins.

## Main

Pooled optical screening is a powerful tool to connect genetic alterations to live-cell phenotypes for libraries of strains (M. Lawson and Elf 2021). In one implementation, the strains in the library are phenotyped under the microscope, without knowledge of their genotype. The genetic identities of the cells are only revealed by *in situ* genotyping after the cells have been fixed (Feldman et al. 2019; Emanuel, Moffitt, and Zhuang 2017; M. J. Lawson et al. 2017; Camsund et al. 2020).

Historically, *in situ* genotyping of bacterial cells has only been possible in cases where the strain-specific phenotypes are induced by a medium to high-copy-number plasmid, ensuring sufficient expression of the genotyping signal. This approach has been used to study the brightness of fluorescent proteins expressed from plasmids (Emanuel, Moffitt, and Zhuang 2017) and to characterise CRISPR knockdown libraries in which a sgRNA is expressed from the same plasmid as the genotyping barcode (Camsund et al. 2020). Many experimental settings do however require the expression of genotyping barcodes from the chromosome. In some experimental designs, variations in the plasmid copy number between different cells could potentially mask differences in the phenotype of interest, as for example when studying noise in gene expression. In other cases, the need for chromosomal expression might be a strict requirement of the experiment to be performed. For example, when studying libraries of fluorescently labelled chromosomal loci, the fluorescent marker itself is integrated together with a barcode to enable efficient identification of the integrated sequence. Similarly, when studying the physiological effects of a mutation in a chromosomal locus, the genotyping barcode would typically be expressed near the locus itself. For the benefit of these experimental designs, we have developed a method for genotyping chromosomal barcodes in *E. coli*.

To demonstrate the use of the method, we chose to characterise the average time it takes from expression to until the protein becomes fluorescent, i.e. the maturation time, of 81 fluorescent proteins (FPs) excitable using 575 nm light. The maturation kinetics of red fluorescent proteins (Balleza, Kim, and Cluzel 2018) is often problematic for bacterial applications, because red FPs often have a longer maturation time than the generation time of many bacteria. In these cases less than half of the FP molecules will be mature at any point in time if expressed under a constitutive promoter. In addition, the slow maturation time of red FPs makes them poorly suitable for studying dynamics of fast cellular processes, as they tend to “blur” the fluorescent readout in time.

We produced a library of red FP genes expressed from the bacterial chromosome. In this case, chromosomal expression normalises the expression levels across strains, and makes the brightness at the selected excitation wavelength directly comparable.

We selected the fluorescent proteins from FPbase (Lambert 2019) and introduced the corresponding *E. coli* codon-optimised open reading frames (ORFs) in the *lacZ* position of the *lac* operon. Each ORF is associated with a unique barcode sequence that is flanked by a T7 promoter (Fig. S1) and stabilising elements to increase the abundance of barcode RNA. Sequences common to all strains are tabulated in Table S1, while the complete gene fragments used to construct each strain are tabulated in Table S2. The strains were constructed individually (see Materials and Methods), but pooled before loading into a mother machine type PDMS microfluidic chip with 4000 cell traps (Baltekin et al. 2017). Each trap fits a single file of 10-16 cells such that after 4 generations, each cell in the trap is a direct descendant of the cell at the bottom and clonally expresses the same FP (Fig. 1A).

**Figure 1:**
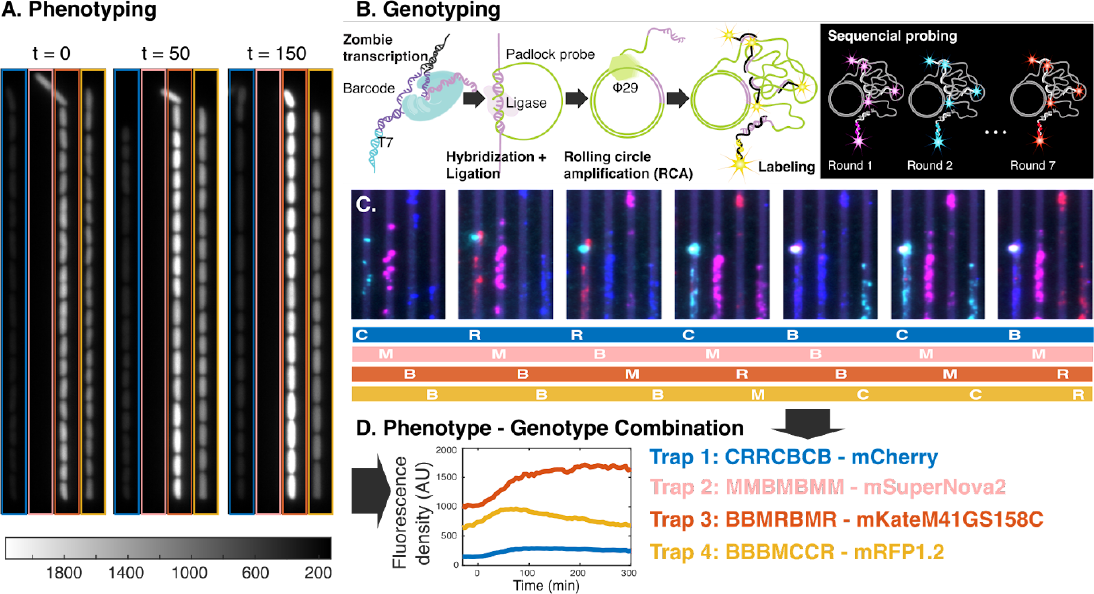
Phenotyping followed by *in situ* genotyping. (A) Example of fluorescence images of cells at three different timepoints after switching to chloramphenicol-containing medium. Time point zero indicates swap to medium with chloramphenicol. (B) Cartoon showing major steps of *in situ* genotyping. (C) Fluorescence images of seven rounds of genotyping for the cell traps shown in A. (D) Quantified fluorescence density throughout the experiment for one example lineage for the traps highlighted in A together with the decoded genotypes from C. None of the cells in Trap 2 passed the filtering criteria on the cell fluorescence and are thus not shown in the plot.

We initiated the maturation experiment with fully induced FP expression at 1 mM IPTG, and imaging in steady-state growth conditions for 1 hour. After this, the media was swapped to media including 250 ug/ml of chloramphenicol, which stops cell growth (Fig. S2) and protein synthesis (Balezza, et al 2018). The already expressed, but still immature, FPs will become fluorescent over time and, in turn, the total cell fluorescence will increase (Fig. 1A & D). The increase of the fluorescence signal was monitored by exposing the cells to 575 nm light every 5 min and detecting the emitted fluorescence. The cell growth was monitored in phase contrast every minute. The experiment was run for a total of 6 hours after which most cells approached a plateau in fluorescence intensity or even showed a fluorescence intensity decrease due to photobleaching.

After collecting the phenotyping data, the genotypes for each cell trap were determined *in situ*. The cells were fixed and permeabilized in 70% EtOH and rehydrated in PBS-Tween before the cell walls were partly degraded by Lysozyme. The RNA barcodes were expressed by adding T7 polymerase (Fig. 1B) (Askary et al. 2020). Barcode-specific ssDNA padlock probes were hybridised directly to the RNA barcodes, ligated by splintR-ligase (Sountoulidis et al. 2020) and amplified by rolling circle amplification (RCA) (Ke et al. 2013). The individual barcodes were read out by sequential FISH hybridization to the RCA products, allowing for detection of a large number of barcodes with a limited set of differently coloured fluorescent probes (Lubeck et al. 2014; K. H. Chen et al. 2015). In each sequential hybridization round, a mix of adaptor “L-probes” were hybridised to the RCA products, followed by parallel hybridization with 4 differently coloured fluorescent detection probes. (See Materials and Methods for details of the genotyping protocol).

For barcode readout, each of the barcodes were assigned a unique sequence of the four differently coloured detection probes, one colour for each of the seven rounds of genotyping. The minimal Hamming distance between the detection probe sequences for the different genotypes was set to 3, allowing correction of a single error in the detection probe sequence (K. H. Chen et al. 2015). See Table 1 for details of the genotyping performances in three replicate experiments. By counting the number of single round errors among the decoded traps we assess an average error probability of 0.041 per trap per round. Assuming that decoding errors are uncorrelated between padlock sequences and rounds of probing, this error probability results in a probability of ∼0.002 for assigning a cell trap with the wrong genotype.

**Table 1:** Genotyping performance in each replicate experiment.

| Exp. id | Nr. of traps with cells | Traps with no signal | Traps with double signal | Unassigned decoded traps | Successfully decoded traps | Decoding error probability |
| --- | --- | --- | --- | --- | --- | --- |
| BY1087 | 3 046 | 893 | 120 | 73 | 1 960 [64.3%] | 0.028 |
| BY1088 | 3 182 | 1 212 | 184 | 177 | 1 609 [50.6%] | 0.068 |
| BY1089 | 3 420 | 1 002 | 212 | 125 | 2 081 [60.8%] | 0.032 |

After decoding the library, we grouped the fluorescence maturation curves based on which FP that was expressed. Solid lines in Figure 2 show the average response for FPs from three replicate experiments. To quantify the maturation time, we fit a single exponential function to the fluorescence intensity integrated over the cell area after background subtraction (Materials and Methods, Fig. 2 dashed lines). For the FPs where the cell fluorescence at the last point of measurement has bleached by more than 15% compared to the peak value (mRFP1.2 and mScarlet-I), a function containing the sum of two exponential functions was used in order to account also the bleaching rate.

**Figure 2.**
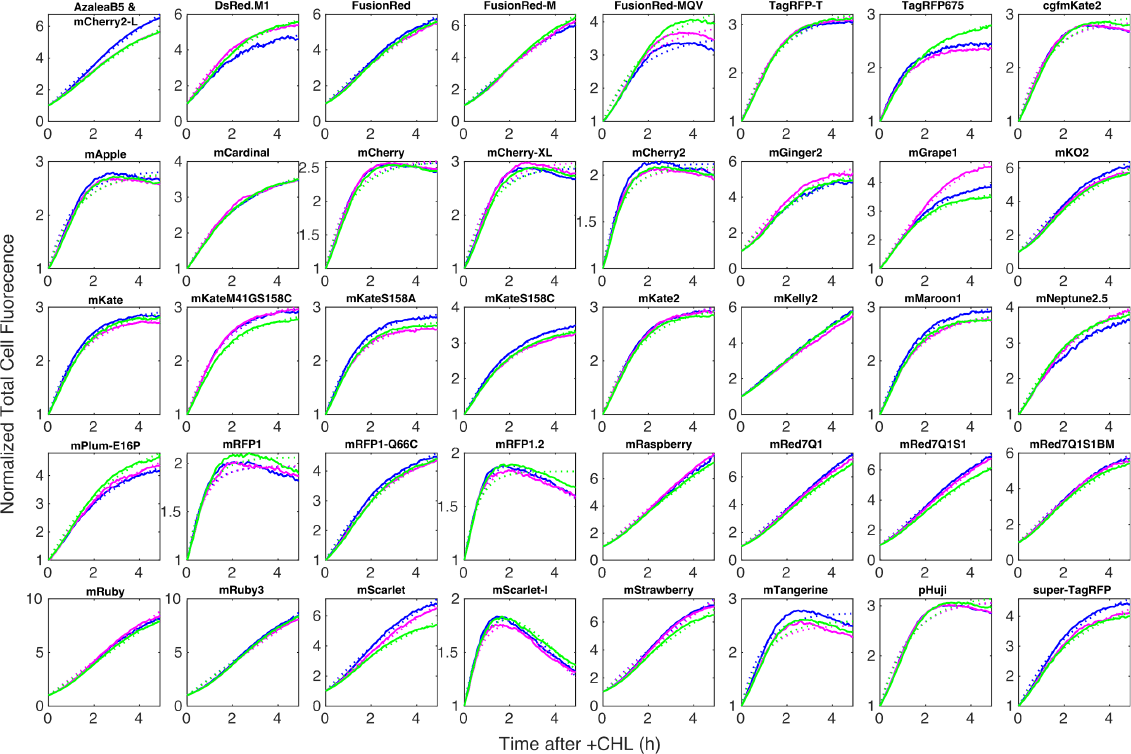
Average total cell fluorescence as function of time after +CHL for different FPs. Different colours represent replicate experiments (blue: BY1087, magenta: BY1088 and green: BY1089). Only FPs where the number of detected cell lineages > 5 for each experiment is shown. Solid lines are experimental data and dotted lines are fits to either a single or double exponential function (see Material and Methods for function definitions).

The maturation times for individual cell lineages together with the average cell-fluorescence intensity in the last three frames of fluorescence imaging before switching to chloramphenicol-containing media are presented in the scatter plots in Figure 3. Note that the brightness is not likely to be the maximal brightness for each fluorophore since all of them are excited using the same light source and the excitation spectra for different fluorophores peaks at different wavelengths. The results from the three replicate experiments are shown in different colours in Figure 3. Single dots in the scatter plots of Figure 3 correspond to measurements from single cell lineages. Statistics of the maturation times shown in Figure 3 are presented in Table S3. When the maturation time is longer than the experiment duration after the switch chloramphenicol-containing medium (300 min), the time constant is reported as >300 min, since these values could not be accurately determined. For all but one FP shown in Figure 3 the majority of the cell lineages fall into one major cluster. The data for barcode 275 is different since both Azaela and mCherry2L FPs mistakenly received the same barcode and this generates two clusters. Using the phenotypic outliers we estimate that ∼3% of the cell traps contain cells with a phenotype that deviate from the majority (Table S4). Since this number is higher than the probability of incorrectly decoding a barcode and genotype calling for many outliers is based on one fluorescent foci, it is most likely due to unspecific padlock ligation.

**Figure 3:**
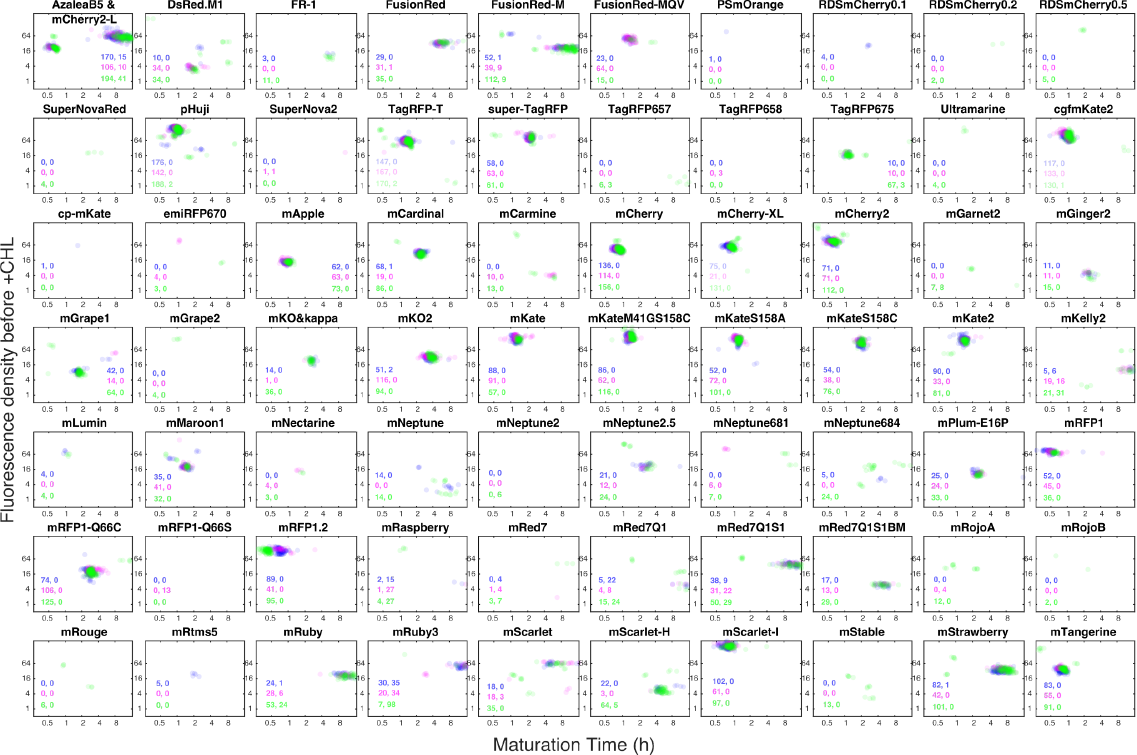
Scatter plots for the maturation time (x-axis) and average cell fluorescence before +CHL (y-axis) for the different FPs. Each dot corresponds to one cell lineage. Colors as in Fig. 2. The number of cell lineages detected in each experiment is reported in the inset using two numbers. The one to the left is the number of data points shown in figure and the number to the right is the number of data points falling outside the axes’ ranges. Each cell trap may contain more than one cell lineage.

We conclude that it is possible to read out single-cell barcodes expressed from the *E. coli* genome in a PDMS microfluidic device. This opens up for large scale genomic engineering and phenotypic analysis with minimal interference from the barcoding system.

## Author contributions

AK, DAGS and BS designed and constructed strains; RS, JL, MG, MN, FS & JE development of genotyping protocol; JL made the experiments; DS and SZ developed barcoding decoding method; DAGS and DF analysed data; J.E & D.F. conceived the project and wrote manuscript with input from all authors.

## Acknowledgments

We are grateful for helpful contributions from Daniel Camsund, Nick Shakari, Nicole Eger and Irmeli Barkefors. This study was made possible by grants from the ERC (advanced grant no. 885360), the Swedish Research Council (grant nos. 2016-06213 and 2018-03958 to JE, as well as 2019-01238 to MN), the Knut and Alice Wallenberg Foundation (grant nos. 2016.0077, 2017.0291, and 2019.0439), and the eSSENCE e-science initiative. The computations and data management were enabled by resources provided by the Swedish National Infrastructure for Computing at UPPMAX, partially funded by the Swedish Research Council through grant agreement no. 2018-05973. J.E. patented optical pooled screening in 2014 (WO2016/007063 A1) and holds shares in Bifrost Biosystems. MN holds shares in Bifrost Biosystems.

## Materials and Methods

### Library construction

*Strain construction and cloning*. We selected 81 fluorescent proteins from the fpbase database (Lambert 2019) with specific properties. Mainly we selected monomeric proteins within the excitation spectrum between 550 - 650 and excluded non-photoswitchable proteins. The amino acid sequence was back translated and codon optimised in Benchling (https://benchling.com (2022)) using *E. coli* as organism and a medium GC content.

To enable chromosomal insertion of these selected fluorescent proteins, we designed gene fragments that incorporated a fluorescent protein under the control of the *lac* promoter and in-frame of the *lacZ* gene (Fig. S1). The terminator between the FP and the BC is a strong synthetic terminator (L3S2P21) and consists of a short RNA hairpin followed by a U-rich sequence (Y.-J. Chen et al. 2013). The barcode is under control of a T7 promoter, and included on the same RNA transcript are two accessory sequences: a spacer and the d0 sequence (Delebecque et al. 2011) that we have used previously because of its enhanced RNA stability (Camsund et al. 2020). As a control we also made a barcoded strain without a FP. For more details on the common motifs see Table S1. 79 gene fragments were ordered from Twist and 3 gene fragments were ordered from IDT. The ordered sequences are in Table S2.

We used lambda red recombination with the selectable/counter selectable marker Atox1 for chromosomal integration (Näsvall 2022). Successful chromosomal integration replaced the Atox1 marker with the wanted construct, leaving no scar or selection marker. We sequenced-verified each strain using Sanger sequencing. The 82 strains were grown overnight on a 96-well plate using 200 μl of LB media, next morning we pooled the strains together to create the redFP-library. The library was stored as a glycerol stock.

### DNA barcode design

We generated *in silico* a set of thousands of random DNA 30-mers, each with a GC content between 40% and 60% and with homopolimeric stretches <4 nucleotides long. We then only selected sequences with a Hamming distance >5 to every other sequence in the dataset. This ensures that the picked sequences are substantially distinct from one another, reducing the probability of spurious detection. From the resulting list, we randomly picked 200 sequences, inserted them into the chromosome at *ygaY* pseudogene with the same design as described above for the library construction. Then we ran an on-chip genotyping experiment to screen for sequences that can be used as barcodes, meaning they are successfully detected using the on-chip genotyping experiment. Essentially all barcodes used in the library were picked from this set of barcodes.

### Padlock probe design

We designed padlock probes (PLPs) against the 82 sequences generated above using a custom script that performs the following steps: each target sequence is reverse-complemented and split in half to produce 5’ and 3’ hybridisation arms. The scaffolds between the 2 arms are then filled with a set of custom sequences, each including a unique readout barcode for sequential hybridisation, a common sequence to prime the RCA reaction, and other accessory sequences that can be used, if necessary, to prime PCR reactions for in-vitro analysis. At the end of this process, each padlock can target one of the 82 sequences above, and can be decoded by a unique sequence of colours across multiple hybridisation cycles. PLPs for the 82 target sequences were pooled at 200nM total concentration and phosphorylated (10ul PLP pool, 2ul T4 PNK (NEB M0201S), 5ul ATP 10mM, 5ul PNK buffer 10x, 28ul nuclease-free water; 30 min @ 37°C, 20 min @ 65°C), resulting in a pool at 0.46nM per oligo. The list of padlock probes used is found in Table S5.

### Phenotyping

#### Media

In the microscopy experiments the media used was M9-succinate (100 μM CaCl_2_, 2 mM MgSO_4_, 1X M9 salts, 0.4% (wt/vol) succinate (Sigma)), 1X RPMI 1640 amino acid mix (Sigma), 0.0425% (wt/vol) Pluronic F108 (Sigma) supplemented with 1 mM IPTG.

#### Optical setup

Microscopy experiments were carried out on a Ti2-E microscope (Nikon) for both phase contrast and epifluorescence. It was equipped with a 100x CFI Plan Apo Lambda DM (Nikon) objective. Images were acquired using a Sona camera (Andor).

In the phenotyping experiment, epifluorescence imaging was carried out using a Spectra III (Lumencor) set to the Yellow channel and a filter cube consisting of a FF01-559/34 (Semrock) excitation filter, a T585lpxr (Chroma) dichroic mirror, and a T590LP (Chroma) emission filter. To maintain a constant temperature of 30ºC, we used a temperature unit and lexan enclosure manufactured by Okolab for the microscope stage and the sample.

In the genotyping experiments, the same microscope setup as for phenotyping was used except for the filter cubes and channel on the Spectra III. The light-source channel and filter combinations used were specific to each dye, as follows: Alexa488: Cyan channel, FF-506-Di03 (Semrock), EX450-490 (Nikon), FF01-524/24 (Semrock); Cy3: Green channel, FF562-Di03(Semrock), FF01-543/22 (Semrock), FF01-586/20 (Semrock); Cy5: Red channel, FF660-Di02 (Semrock), 692/40 (Semrock); Alexa750: NIR channel, T760lpxr (Chroma), ET811/80 (Chroma).

#### The microfluidic chip

The chip is made of PDMS (SYLGARD 184) and it is bonded to a glass No. 1.5 coverslip (Menzel-Gläser) (see (Baltekin et al. 2017) for details). Flow direction and rate was controlled by an OB1 MkIII (Elveflow) electro pneumatic controller.

#### Cell growth before loading in the microfluidic chip

The pooled library was grown overnight in experiment media at 37ºC in a shaking incubator. Next day, cells were diluted 1:250 in fresh media and grown for 4 hours before loading.

#### Maturation time measurements

After loading into the microfluidic chip, cells were grown for about 4 hours. Either 80 (BY1087) or 100 (BY1088 and BY1089) positions were imaged for 6 hours every minute in phase contrast (80 ms exposure time) and every 5 minutes in epifluorescence (100 ms exposure time). 1 hour after the imaging was started the media was swapped to a one containing chloramphenicol (at 250 μg/ml).

### Genotyping

After phenotyping, cells were fixed and permeabilized with 70% EtOH for 20 minutes followed by rehydration in PBS 0.1% tween (PBS-T) overnight in RT.

From this stage all reaction mixes are administered to the front channel of the chip using a syringe pump and the lexan stage enclosure is kept at 30 degrees. The composition of the reaction mixes are given in Table S6 and are administered as described below: 1) To further improve cell permeabilization we flow the lysozyme reaction mix at 1 μl/min for 10 mins onto the chip; 2) Lysozyme activity was stopped by flowing the BSA solution mix at 1ul/min for 10 mins; 3) To produce the RNA barcodes used for Padlock probe binding we flowed the zombie transcription mixture to the chip at a rate of 0.5 μl/min for 120 min; 4) To allow the padlock probe hybridization to the corresponding RNA target we flow the padlock probe hybridization mixture to the chip at 1μl/min for 30 min; 5) For the ligation of the Padlocks hybridised to the target RNA, a SplintR ligase mixture was flown into the chip at 0.75 μl/min for 60 min; 6) For amplification of the Padlock DNA we flow the RCA reaction mixture onto the chip at 0.5 μl/min for 120 min. 7) Barcode detection: for each round of sequential FISH, the L-probes, pre-hybridised to detection oligos, were flown onto the chip at 1 μl/min for 30 min followed by imaging; 8) Probes are removed from the RCA product by flowing the probe stripping buffer at 1 μl/min for 30 min. Step 7 and 8 were repeated an additional 6 times with different variations of L-probe detection oligo pools.

The L-probes were mixed for each hybridization-round according to the code allowing for error correction (See Table S7 for a list of the L-probes used). Each strain was assigned a randomly chosen 7-symbol code word (with an alphabet of 4 symbols), while ensuring that each code word has a Hamming distance of at least 3 symbols from all other 81 code words. Each symbol in the code is represented by a differently colored detection oligo. Thus, for each hybridization-round we mixed a collection of 82 L-probes each matching the corresponding detection oligo per strain.

### Analysis

#### Image analysis

Image analysis was done using an in-house pipeline. In the pipeline, cells are first segmented using a U-Net convolutional network based approach (Ronneberger, Fischer, and Brox 2015). The segmented cells are then tracked from frame to frame and linked into lineages using the Baxter algorithm (Magnusson et al. 2015). Total cell fluorescence was calculated by summing up camera pixel intensities inside the segmented cell outline. To only include pixel intensity due to fluorescence, the average pixel intensity of an area without either cells or PDMS bonded directly to glass, was subtracted from each pixel value before making the sum in each cell. Cells without an FP showed no discernible signal. Given the high fluorescence intensity variability between different FPs and the small spatial distance between adjacent cell traps in the microfluidic chip, the signal from cells carrying a high intensity FP may overshadow the signal of cells in neighbouring traps which carry a low intensity FP. To overcome the problem with signal bleeding between cell traps, we only included cells where the intensity density inside the segmented cells is, on average, 3-fold higher as compared to the cell’s immediate surroundings. The tracked cell lineages were used to generate one maturation curve for each entire lineage where the areas and total fluorescence of two daughters, four granddaughters, etc, were summed up for each time point. Only cell lineages that existed at the time of swapping to chloramphenicol-containing medium and that were possible to track until the end of the experiment were used in calculating the maturation times.

#### Maturation time calculations

To estimate the FP maturation time, the total cell fluorescence after chloramphenicol treatment is fitted to either a function consisting of a single exponential term, or a function consisting of the sum of two exponential terms. These model fits assumes that the maturation process is well described by a single step reaction and that protein production is instantly, and completely, stopped at the time of exposing the cells to chloramphenicol. Residual expression of the fluorescent protein after the switch to chloramphenicol-containing medium may skew the maturation time estimates, but will not affect the maturation time rank order.

In cases where where the total fluorescence is reaching a constant plateau, it is fitted to the function

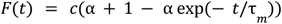

where *F* is the total cell fluorescence, *t* is time after chloramphenicol addition, *τ*_*m*_ is the maturation time, and *c* and *α* are fitting constants.

In cases where the total fluorescence is clearly decreasing at the end of the experiment such that the ratio between the peak fluorescence and the end point fluorescence, on average for all cells in one experiment carrying the same FP, is less than 0.85, the total fluorescence after chloramphenicol addition is fitted to

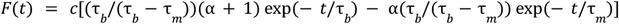

where *τ*_*b*_ is the characteristic time for signal decrease due to bleaching.

#### Phenotypic outlier detection

To quantify the number of cell traps which are showing a phenotype that is deviating from the behaviour of the majority of cells in traps genotyped as containing the same FP, we used a clustering based analysis of the data shown in Fig. 3. First the data from the three different repeat experiments were pooled. After this, we used DBSCAN to find clusters. The cluster containing the largest number of cells was identified as the major cluster. Finally the number of traps in the major cluster and the number of traps outside the major cluster were counted. Only traps where the number of identified lineages were more than 2, and where all the cells in the trap were unanimously identified as either being inside or outside the major cluster were used. Only FPs where the number of traps in the major cluster where more than 5 were included. The fraction of traps with deviating phenotypes are shown in Table S4.

#### Barcode detection

Phase contrast images from each round of genotyping were aligned using the dot barcodes imprinted next to empty traps. Trap segmentation was performed using the deep learning model described in (Kandavalli et al. 2022). Empty traps were detected by differentiating between empty traps that have a smooth image, compared to traps full with lysed cells that result in a rugged image. This was done by thresholding the magnitude of the highest bins in a Fourier transform of the cropped traps images. Fluorescent blobs were detected using a signal threshold that was four standard deviations higher than the mean of the Gaussian fitted to the histogram of pixel intensities from all of the traps in each position in the same fluorescence channel. The resulting fluorescence masks were compared between rounds and the final mask comprised of the pixels that appear at least in half of the rounds. For trap code assignment, if the signal within the fluorescence mask for one channel was at least twice as strong than for the other channels it was assigned as the true channel for that round. We applied error correction to trap codes fixing at most 1 round, by assigning the decoded sequence to one of the code words if it was identical to this code word, or one error away from it (*i*.*e*. Hamming distance equals 1).

