## Supplementary Figures and Tables for "Pooled optical screening in bacteria using chromosomally expressed barcodes"

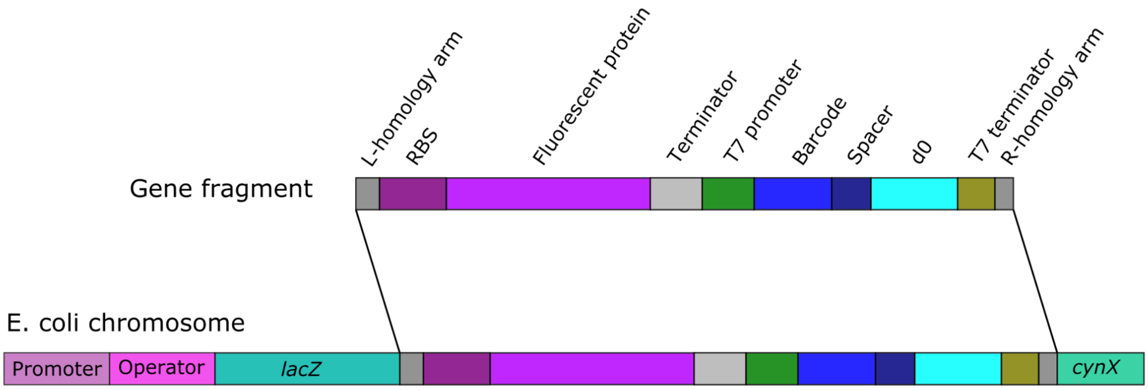

**Figure S1. Diagram of design**

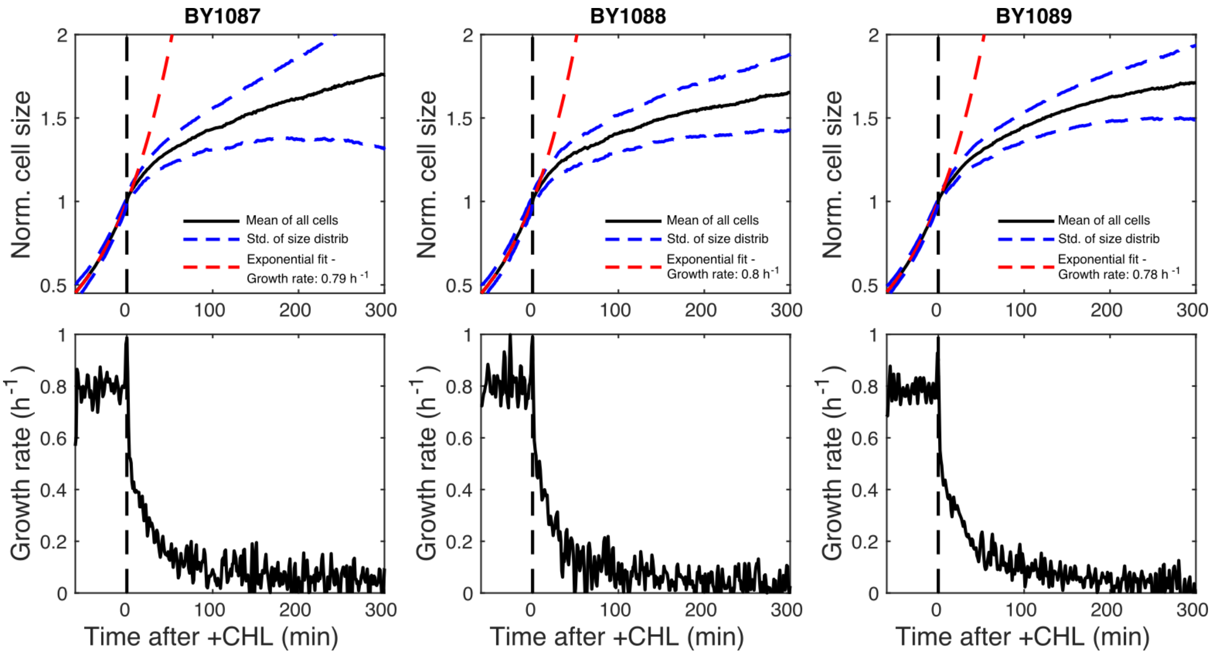

**Figure S2: Cell size and growth rate change as response to +CHL.** Cell size as function of time (top row) and growth rate as function of time (bottom row) for three replicate experiments. Black solid line is the average of all cells in each experiment. Dash blue curves indicate one standard deviation for the distribution of cell sizes for each time point. Red dashed line shows regression to single exponential before swap to +CHL. Fitted growth rate shown in inset.

### Supplementary Tables

| **Motif** | **DNA sequence** |
| --- | --- |
| R-homology arm | AATCCGCCGTTTGTTCCCACGGAGAATCCGACGGGTTGTTACTCGCTCACATTTAATGTT |
| FP-RBS | TTTGTTTAACTTTAAGAAGGAGA |
| terminator (L3S2P21) | CTCGGTACCAAATTCCAGAAAAGAGGCCTCCCGAAAGGGGGGCCTTTTTTCGTTTTGGTCC |
| T7-promoter | TAATACGACTCACTATAGGGAGA |
| Spacer | GGTTGACCTTTGTACATTAATTTAA |
| d0 | GGGAGGACTCCCACAGTCACTGGGGAGTCCTCGAATACGAGCTGGGCACAGAAGATATGGCTT CGTGCCCAGGAAGTGTTCGCACTTCTCTCGTATTCGATTCCC |
| T7 terminator | CTAGCATAACCCCTTGGGGCCTCTAAACGGGTCTTGAGGGGTTTTTTG |
| L-homology arm | CATATCGAATTTACGGCTAGCTCAGTCCTAGGTATAGTGCTAGCGCAAGGAGACAAGAGA |

**Table S1: DNA sequences of common motifs**

**Table S2: DNA sequences for complete chromosome insert for all strains.**

| | | BY1087 | | BY1088 | | BY1089 | |
|---|---|---|---|---|---|---|---|
| BC | FP | median (min) | nr of lineages | median (min) | nr of lineages | median (min) | nr of lineages |
| 275 | AzaleaB5 | >300 | 185 | >300 | 116 | >300 | 235 |
| 280 | DsRed.M1 | | 0 | 112 | 34 | 130 | 34 |
| 283 | FusionRed | >300 | 29 | >300 | 32 | >300 | 35 |
| 284 | FusionRed-M | >300 | 53 | >300 | 48 | >300 | 121 |
| 285 | FusionRed-MQV | 77 | 23 | 81 | 64 | 91 | 15 |
| 301 | pHuji | 57 | 176 | 59 | 142 | 62 | 190 |
| 308 | TagRFP-T | 78 | 147 | 81 | 167 | 88 | 172 |
| 309 | super-TagRFP | 131 | 58 | 135 | 63 | 141 | 61 |
| 312 | TagRFP675 | | 0 | | 0 | 71 | 70 |
| 317 | cgfmKate2 | 58 | 117 | 60 | 133 | 66 | 131 |
| 328 | mApple | 55 | 62 | 57 | 63 | 62 | 73 |
| 330 | mCardinal | 141 | 69 | 135 | 19 | 144 | 86 |
| 335 | mCherry | 48 | 136 | 50 | 114 | 54 | 156 |
| 337 | mCherry-XL | 47 | 75 | 53 | 21 | 58 | 131 |
| 338 | mCherry2 | 34 | 71 | 35 | 71 | 39 | 112 |
| 356 | mGinger2 | | 0 | | 0 | 155 | 15 |
| 357 | mGrape1 | 101 | 42 | | 0 | 108 | 64 |
| 363 | mKO&kappa | | 0 | | 0 | 155 | 36 |
| 367 | mKO2 | 205 | 53 | 208 | 116 | 232 | 94 |
| 369 | mKate | 74 | 88 | 75 | 91 | 83 | 57 |
| 370 | mKateM41GS158C | 80 | 86 | 84 | 62 | 90 | 116 |
| 371 | mKateS158A | 73 | 52 | 72 | 72 | 75 | 101 |
| 372 | mKateS158C | 116 | 54 | 114 | 38 | 121 | 76 |
| 373 | mKate2 | 82 | 90 | 83 | 33 | 87 | 81 |
| 378 | mKelly2 | | 0 | >300 | 35 | >300 | 52 |
| 382 | mMaroon1 | 81 | 35 | 85 | 41 | 89 | 32 |
| 388 | mNeptune2.5 | 149 | 21 | | 0 | 169 | 24 |
| 395 | mNeptune684 | | 0 | | 0 | 186 | 24 |
| 403 | mPlum-E16P | 141 | 25 | 159 | 24 | 151 | 33 |
| 404 | mRFP1 | 29 | 52 | 31 | 45 | 35 | 36 |
| 409 | mRFP1-Q66C | 163 | 74 | 177 | 106 | 178 | 125 |
| 418 | mRFP1.2 | 43 | 89 | 43 | 41 | 25 | 95 |
| 424 | mRaspberry | | 0 | >300 | 28 | >300 | 31 |
| 426 | mRed7Q1 | >300 | 27 | | 0 | >300 | 39 |
| 429 | mRed7Q1S1 | >300 | 47 | >300 | 53 | >300 | 79 |
| 430 | mRed7Q1S1BM | 298 | 17 | | 0 | 285 | 29 |
| 436 | mRuby | >300 | 25 | >300 | 34 | >300 | 77 |
| 438 | mRuby3 | >300 | 65 | >300 | 54 | >300 | 105 |
| 450 | mScarlet | >300 | 18 | >300 | 21 | >300 | 35 |
| 454 | mScarlet-H | 284 | 22 | | 0 | >300 | 69 |
| 457 | mScarlet-I | 47 | 102 | 50 | 61 | 53 | 97 |
| 464 | mStrawberry | >300 | 83 | >300 | 42 | >300 | 101 |
| 468 | mTangerine | 45 | 83 | 45 | 55 | 52 | 91 |

**Table S3. Maturation time summary**

| Experiment | Nr. of outlier traps | Total nr. of traps | Fraction |
|---|---|---|---|
| BY1087 | 3 | 351 | 0.009 |
| BY1088 | 4 | 288 | 0.014 |
| BY1088 | 25 | 498 | 0.050 |
| Total | 32 | 1137 | 0.028 |

**Table S4: Frequencies of cell traps containing cells with phenotypic outliers.**

**Table S5: Padlock probe sequences**

| **1. Lysozyme reaction mix** | **Stock Conc.** | **Final Conc.** | **Volume (uL)** |
|---|---|---|---|
| Lysozyme (Thermo scientific) | 50 mg/ml. | 250 ug/ml | |
| PBS-Tween 0,01% | | | |
| **2. BSA solution** | **Stock Conc.** | **Final Conc.** | **Volume (uL)** |
| BSA (Thermo scientific) | 10% | 1% | |
| PBS-Tween 0,01% | | | |
| **3. Zombie transcription mixture** | **Stock Conc.** | **Final Conc.** | **Volume (uL)** |
| Transcription buffer (Thermo scientific) | 5x | 1x | 16 |
| NTPs | 25 mM | 2 mM | 6.4 |
| MgCl2 | 25 mM | 6 mM | 19.2 |
| Tween 20 (Promega) | 1% (v/v) | 0.1% (v/v) | 8 |
| Glycerol | 50% | 5% | 8 |
| T7 RNA Polymerase (Thermo scientific) | 20 U/µl | 2 U/µl | 8 |
| Riboprotect (Qiagen gdansk) | 40 U/µl | 1 U/µl | 2 |
| H2O mQ | - | - | 12.4 |
| **4. PLP Hybridization mixture** | **Stock Conc.** | **Final Conc.** | **Volume (uL)** |
| 20x SSC | 20x | 2x | 5 |
| Ethylene Carbonate (Merch) | 100% | 5% | 2.5 |
| MgCl2 | 50 mM | 15 mM | 15 |
| Tween 20 | 1% (v/v) | 0.1% (v/v) | 5 |
| Padlock probe library (each probe) | 0.46 nM | 0.2 nM | 21.25 |
| Riboprotect (Qiagen gdansk) | 40 U/µl | 1 U/µl | 1.25 |
| **5. SplintR ligation mixture** | **Stock Conc.** | **Final Conc.** | **Volume (uL)** |
| SplinR Buffer (NEB) | 10x | 1x | 6 |
| Glycerol | 50% | 5% | 6 |
| Tween 20 | 1% (v/v) | 0.1% (v/v) | 6 |
| Riboprotect (Qiagen gdansk) | 40 U/µl | 1 U/µl | 1.5 |

| | | | |
|---|---|---|---|
| SplintR Ligase (NEB) | 25 U/uL | 0.5 U/uL | 1.2 |
| H2O mQ | - | - | 39.3 |
| **6. RCA mixture** | **Stock Conc.** | **Final Conc.** | **Volume (uL)** |
| dNTPs | 2.5 mM | 250 uM | 9 |
| phi 29 buffer * | 10x | 1 x | 9 |
| Glycerol | 50% | 5% | 9 |
| BSA | 20 µg/µl | 0.2 µg/µl | 0.9 |
| phi 29 polymerase * | 10 U/uL | 1 U/uL | 9 |
| H2O mQ | - | - | 53.1 |
| **7. Labeling mixture** | **Stock Conc.** | **Final Conc.** | **Volume (uL)** |
| Detection oligos (IDT) | 1 µM each | 100 nM | 6 |
| 4x SSC 40% Formamide | 2x | 1x | 30 |
| L-probe pool | 1 nM total | 0.1 nM total | 6 |
| H2O mQ | - | - | 18 |
| **8. Probe stripping mixture** | **Stock Conc.** | **Final Conc.** | **Volume (uL)** |
| Formamide | 100% | 90% | 900 |
| PBS-Tween | 1x | 1x | 100 |

**Table S6: Reaction mixes used for *in situ* genotyping.** *Wild-type phi29 DNA polymerase was transformed into *E. coli* BL21 (DE3) T1R pRARE2 cells. The cells were cultivated in Terrific Broth (TB) medium. Protein expression was induced with IPTG and the protein was purified by immobilized metal-ion chromatography (IMAC), followed by size exclusion chromatography (SEC). The phi29 reaction buffer at 10x concentration is: 500 mM Tris-HCl (pH 8.3), 100 mM $MgCl_2$ & 100 mM $(NH_4)_2SO_4$

**Table S7: L-probe sequences**
